# Bayesian non-central chi regression for neuroimaging

**DOI:** 10.1101/095844

**Authors:** Bertil Wegmann, Anders Eklund, Mattias Villani

## Abstract

We propose a regression model for non-central *χ* (NC-*χ*) distributed functional magnetic resonance imaging (fMRI) and diffusion weighted imaging (DWI) data, with the heteroscedastic Rician regression model as a prominent special case. The model allows both parameters in the NC-*χ* distribution to be linked to explanatory variables, with the relevant covariates automatically chosen by Bayesian variable selection. A highly efficient Markov chain Monte Carlo (MCMC) algorithm is proposed for simulating from the joint Bayesian posterior distribution of all model parameters and the binary covariate selection indicators. Simulated fMRI data is used to demonstrate that the Rician model is able to localize brain activity much more accurately than the traditionally used Gaussian model at low signal-to-noise ratios. Using a diffusion dataset from the Human Connectome Project, it is also shown that the commonly used approximate Gaussian noise model underestimates the mean diffusivity (MD) and the fractional anisotropy (FA) in the single-diffusion tensor model compared to the theoretically correct Rician model.

## 1. Introduction

Gaussian statistical models are very common in the field of neuroimaging, as they enable efficient algorithms for estimation of brain activity and connectivity. However, the measured signal in diffusion weighted imaging (DWI) and functional magnetic resonance imaging (fMRI) is the magnitude of a complex-valued Gaussian signal and therefore follows a Rician distribution, see Gudbjartsson and Patz (1995) and Section 2.1. The Gaussian model is a good approximation to the Rician model in fMRI as the signal-to-noise (SNR), defined here as the ratio of the average BOLD signal to its standard deviation, for fMRI data tends to be large enough for the approximation to be accurate (Adrian et al., 2013). However, the recent push towards higher temporal and spatial resolution in neuroimaging (Moeller et al., 2010; Feinberg and Yacoub, 2012; Setsompop et al., 2013) may lead to low SNRs with increased risk of distorted conclusions about brain activity and connectivity. This is demonstrated in Section 4, where a Rician model is able to accurately detect brain activity at low SNRs, while the Gaussian approach fails to do so. Low SNRs are also common for DWI, especially when the b-value is high (Zhu et al., 2009). Using a Gaussian model for diffusion tensor imaging (DTI) can therefore lead to severely misleading inferences. The reason for the popularity of the Gaussian approach is that Gaussian models can be analyzed using simple algorithms, while the Rician distribution is complicated since it does not belong to the exponential family. More generally, MR images collected by simultaneous acquisition from L independent coils may follow the non-central *χ* (NC-*χ*) distribution with L degrees of freedom, depending on how the measurements are combined into a single image (Tristán-Vega et al., 2012; Aja-Fernandez and Vegas-Sanchez-Ferrero, 2016). We therefore derive our algorithm for the general NC-*χ* model from which the Rician model can be directly obtained as the special case when *L* = 1.

### 1.1. Rician models in fMRI

There have been a handful of approaches for the Rician model in fMRI applications. Solo and Noh (2007) and Zhu et al. (2009) propose to augment each data observation with the missing phase information, and to use the EM algorithm to obtain the maximum likelihood estimates of the regression coefficients in the Rician model; Adrian et al. (2013) provide the extension to the case with autocorrelated errors. Although not discussed in the literature, the data augmention technique is naturally extended to a fully Bayesian analysis via Gibbs sampling, where the parameters are iteratively sampled conditional on the missing phase observations, followed by a sampling step for the phases given the model parameters. The convenience of introducing unobserved phase information does not come without cost, however, and data augmentation is well known to lead to inefficient exploration of the posterior distribution and inflated numerical standard errors (Liu et al., 1994). The same problems tend to plague the EM algorithm, which often exhibit very slow convergence.

### 1.2. Rician models in DTI

Rician models have mainly been used for noise removal in DTI (Basu et al., 2006; Wiest-Daesslé et al., 2008; Aja-Fernandez et al., 2008), but also for tensor estimation (Andersson, 2008; Veraart et al., 2011). The only method that we are aware of for estimating diffusion parameters in the more general NC-*χ* regression model, for data acquired with several independent coils, is the (non-Bayesian) least squares approach presented by Tristán-Vega et al. (2012).

### 1.3. Non-central chi regression

We therefore introduce a NC-*χ* regression model where both parameters in the distribution (the mean and variance of the underlying complex-valued signal) are modeled as functions of covariates, with the Rician model as an important special case. We propose a Bayesian analysis of the model based on a highly efficient Markov Chain Monte Carlo (MCMC) algorithm, to simulate from the joint posterior distribution of all model parameters. Contrary to previous Bayesian and EM approach, our Bayesian methods works directly on NC-*χ* or Rician distributions, without the need to introduce missing phase data, and the MCMC convergence is excellent due to an accurately tailored proposal distribution. A high efficiency makes it possible to use a smaller number of simulations to obtain the same numerical accuracy. This is absolutely crucial for imaging applications since a separate MCMC chain is run for each voxel. Moreover, our MCMC algorithm also performs Bayesian variable selection among both sets of covariates. For both DTI and fMRI data, our Bayesian approach has the obvious advantage of capturing the uncertainty in each voxel. The uncertainty can easily be propagated to the group analysis, to down-weight subjects with a higher uncertainty. This is in contrast to the popular TBSS approach (tract-based spatial statistics) (Smith et al., 2006) for voxel-wise multi-subject analysis of fractional anisotropy (FA), which ignores the uncertainty of the FA.

Using a freely available DWI dataset from the Human Connectome Project (Essen et al., 2013), we show that commonly used Gaussian DTI approximation underestimates the mean diffusivity (MD) and substantially underestimates the FA of the single-diffusion tensors, compared to the theoretically motivated Rician model, especially in white-matter regions with high FA. In addition, we show that covariates are needed in both parameters of the Rician distribution, not only in the mean. In an fMRI simulation study, we formulate a sensible prior distribution for the regression coefficients based on the Fisher information matrix, and demonstrate that the Rician model is remarkably adept at recovering the activations even at very low SNRs. We also show that the Gaussian model is likely to lead to severely erroneous activation inference in such settings.

### 1.4. Application to more advanced diffusion models

We have here focused on the rather simple single-diffusion tensor, while more recent work focus on extending the diffusion tensor to higher orders. In the work by Westin et al. (2016), a regression approach is used to estimate the diffusion tensor and a fourth order covariance matrix in every voxel. Our regression framework can therefore easily be applied to QTI (q-space trajectory imaging) data (Westin et al., 2016) as well, and more generally for any diffusion model that can be estimated using regression. Moreover, DTI is still the most common choice for studies investigating FA differences between healthy controls and subjects with some disease (Shenton et al., 2012; Eierud et al., 2014).

## 2. Heteroscedastic rician and NC-*χ* regression

We start by describing our model for the special case of a Rician distribution, and then generalize it to the NC-*χ* case.

### 2.1. Rician regression

The measured signal in DTI and fMRI is a complex-valued indirect measure of structural brain connectivity and brain activity, respectively,

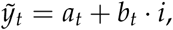

where the real part *a_t_* ~ *N* (*μ_t_* cos *θ_t_*, *ϕ_t_*) and the imaginary part *b_t_* ~ *N* (*μ_t_* sin *θ_t_*, *ϕ_t_*) are independent, and the mean

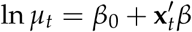

is a linear function of a vector of covariates **x***_t_* at measurement *t*. In fMRI the vector **x***_t_* typically contains the stimulus of the experiment convolved with a hemodynamic response function, polynomial time trends and head motion parameters, while **x***_t_* mainly contains gradient directions in DTI. Note that *ϕ_t_* is potentially measurement-varying, to allow for heteroscedastic complex-valued noise.

It is rare to analyze the complex signal measurements *a_t_* and *b_t_* directly ((Rowe and Logan, 2004) and follow-up papers are exceptions). The most common approach is to use the magnitude of *ỹ_t_* as response variable, i.e.

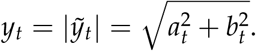

It is well-known that the magnitude follows a Rician distribution (Rice, 1945) with density function

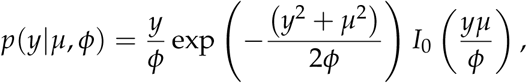

for *y* > 0 and zero otherwise.

The discussion above uses *t*, as in time, as subscripts for the observations. This is suitable for fMRI time series, but to emphasize that our models can also be used for DWI data (see Section 5.1), we will in the remainder of the paper use the more generic *i* to denote observations. We propose the following heteroscedastic Rician regression model

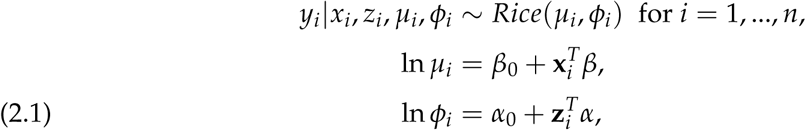

and independence of the *y_i_ conditional* on the covariates in **x***_i_* and **z***_i_*. Since **x***_i_* and **z***_i_* may contain lags of the response variables, our model can capture temporal dependence in fMRI. Note also that we allow for heteroscedasticity in the complex signal, since the variance of the underlying complex-valued signal *ϕ_i_* is a function of the regressors in **z***_i_*. Although the model in Eq. 2.1 has the same structure as a generalized linear model (GLM) (McCullagh and Nelder, 1989of squared magnitudes follow the), it is actually outside the GLM class since the Rician distribution does not belong to the exponential family. The logarithmic link functions used in Eq. 2.1 can be replaced by any twice-differentiable invertible link function.

### 2.2. NC-*χ* regression

Both fMRI and DWI images may be obtained from parallel acquisition protocols with multiple coils, often used to increase the temporal and spatial resolution. Under the assumption of independent complex Gaussian distributed noise in each coil, the sum of squared magnitudes follow the non-central *χ* (NC-*χ*) distribution (Tristán-Vega et al., 2012; Aja-Fernandez and Vegas-Sanchez-Ferrero, 2016). The non-central *χ* density with 2*L* degrees of freedom is of the form

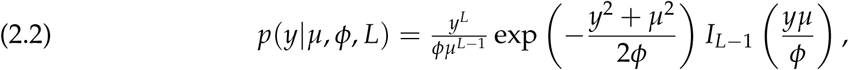

for *y*, *μ*, *ϕ* > 0. We denote this as *y* ~NC-*χ*. Note that when *L* = 1, the density in Eq. 2.2 reduces to the Rice (*μ*, *ϕ*) density. Similarly to the Rician case, we can model *μ* and *ϕ* as functions of explanatory variables via logarithmic link functions. In summary, the observations are assumed to be independently NC-*χ* distributed conditional on the explanatory variables, according to

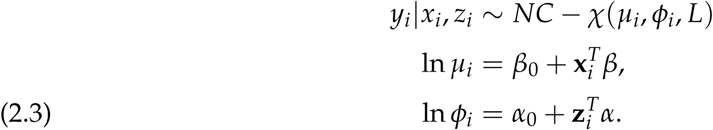

Lagged response values may again be used as covariates in *μ* and *ϕ* to induce temporal dependence.

The order *L* of the NC-*χ* distribution may be given by the problem at hand, for example by the number of independent coils used for data collection. Due to the lack of perfect independence between coils and other imperfections, *L* is often unknown and needs to be estimated from the data. Note that *L* can in general be any positive real number in the NC-*χ* distribution, and does not need to be an integer. Our approach makes it straightforward to introduce an MCMC updating step, to simulate from the conditional posterior distribution of ln *L*, or even model ln *L* as a linear function of covariates.

## 3. Bayesian inference

The Bayesian approach formulates a prior distribution for all model parameters, and then updates this prior distribution with observed data through the likelihood function to a posterior distribution.

### 3.1. Posterior distribution and posterior probability maps

The aim of a Bayesian analysis is the joint posterior distribution of all model parameters

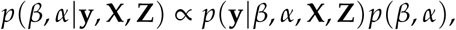

where *p*(**y**| *β*,*α*,**X**,**Z**) is the likelihood function for the MR signal, *p*(*β*,*α*) is the prior, 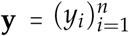,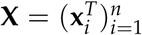 and 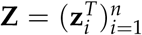; we are here including the intercepts in *β* and *α* Based on this joint posterior one can compute the marginal posterior of any quantity of interest. From the joint posterior *p*(*β*,*α* |**y**, **x**, **z**), it is straight forward to compute Posterior Probability Maps (PPMs), see Friston and Penny (2003). For fMRI, the PPM is an image of the marginal posterior probabilities of positive activation, Pr(*β_j_* > 0|**y**, **X**,**Z**), if the predicted BOLD is the jth covariate in **x**. The joint posterior *p*(*β*,*α*|**y**,**X**,**Z**) for the Rician and NC-*χ* regression models is intractable, and we instead simulate from the joint posterior using an efficient MCMC algorithm described in Section 3.4.

### 3.2. Prior distribution

Our prior distribution for the Rician and the NC-*χ* model is from the general class in Villani et al. (2012). Let us for clarity focus on the prior for *β*_0_ and *β* in ln 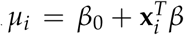; the prior on *α*_0_ and *α* in *ϕ* is completely analogous. We first discuss the prior on the intercept *β*_0_. Start by standardizing the covariates to have mean zero and unit standard deviation. This makes it reasonable to assume prior independence between *β*_0_ and *β*. The intercept is then ln *μ* at the mean of the original covariates. The idea is to let the user specify a prior directly on *μ* when the covariates are at their means, and then back out the implied prior on *β*_0_. Let *μ* have a log-normal density with mean *m*\* and variance *s*\*^2^.

The induced prior on the intercept is then *β*_0_ ~ *N*(*m*,*s*^2^) with 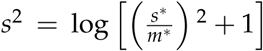 and *m* = log(*m*\*) — *s*^2^/2.

The prior on *β* needs some care, since its effect on the response comes through a link function, and *μ* enters the model partly via a non-linear Bessel function. Following Villani et al. (2012), we let *β* ~ *N*(0,*c*Σ), where Σ = (*X^T^D̂X*)^−1^ is the Fisher information for *β*, *X* is the matrix of covariates excluding the intercept, and *D̂* is the Fisher information for *μ* conditional on *ϕ*, evaluated at the prior modes of *β*_0_ and *β*, i.e. at the vector (*m*, 0^′^)^′^. Thus *D̂* depends only on the constant *m*. The conditional Fisher information for *μ* = (*μ*_1_,… *μ*_n_)′ is a diagonal matrix with elements

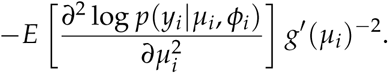

Setting *c* = *n* gives a unit information prior, i.e. a weak prior that carries the information equivalent to a single observation from the model.

### 3.3. Variable selection

Our MCMC algorithm can perform Bayesian variable selection among both sets of covariates (i.e. **x** and **z**). We make the assumption that the intercepts in ln *μ* and ln *ϕ* are always included in the model. Let us again focus on *β* in the equation for *μ*. Define the vector with binary indicators 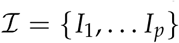 such that *I_j_* = 0 means that the *j*th element in *β* is zero, and that the corresponding covariate drops out of the model. Let 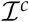 denote the complement of 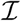. Let 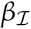 denote the subset of regression coefficients selected by 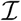. To allow for variable selection we take the previous prior *β* ~ *N*(0, *c*Σ) and condition on the zeros in *β* dictated by 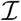 :

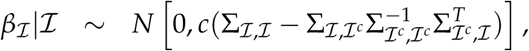

and 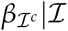 is identically zero. To complete the variable selection prior, we let the elements of 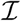 to be a priori independent and Bernoulli distributed, i.e.Pr(*I_i_* = 1) = *π*, and *π* is allowed to be different for the covariates in *μ* and *ϕ*. We choose *π* = 0.5 for both sets of covariates in *μ* and *ϕ*. Other priors on 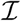 are just as easily handled.

### 3.4. Markov chain monte carlo algorithm

We use the Metropolis-within-Gibbs sampler presented in Villani et al. (2009) and Villani et al. (2012). The algorithm samples iteratively from the set of full conditional posteriors, which in our case here are

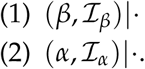

Note that we sample *β* and 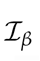 jointly given the other parameters (indicated by ·). The full conditional posteriors 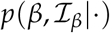 and 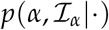 are highly non-standard distributions, but can be efficiently sampled using tailored Metropolis-Hastings (MH) updates. The sampling of the pair 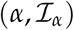 is analoguous to the sampling of 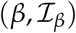, so we will only describe the update of 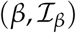. The MH proposal distribution is of the form

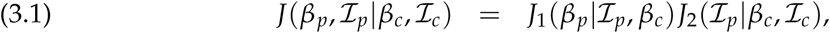

where 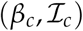denotes the current and 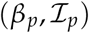 the proposed posterior draw. Following Villani et al. (2009), we choose *J*_2_ to be a simple proposal of 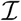 where a subset of the indicators is randomly selected and a change of the selected indicators is proposed, one variable at a time. The proposal of *β*, the *J*_1_ distribution, is a multivariate-*t* distribution with *ν* degrees of freedom:

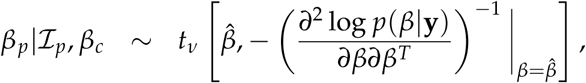

where 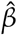 is the terminal point of a small number of Newton iterations to climb towards the mode of the full conditional 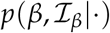, and 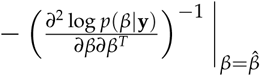 is the negative inverse Hessian of the full conditional posterior evaluated at 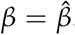 Note that we are for notational simplicity suppressing the conditioning on the covariates **X** and **Z**.

There are a number of different aspects of these Newton-based proposals. First, the number of Newton iterations can be kept very small (one or two steps is often sufficient), since the iterations always start at *β_c_*, which is typically not far from the mode. Second, 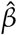 is often not exactly the mode, but the posterior draws from the algorithm will nevertheless converge to the underlying target posterior. Third, the update 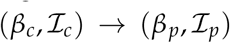 is accepted with probability

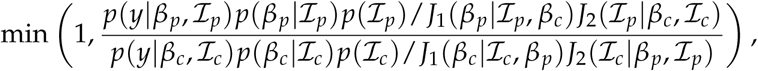

where the factor *J*_1_ 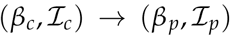 is computed from another round of Newton iterations, this time starting from the proposed point *β_p_*. Fourth, to implement the Newton iterations we need to be able to compute the gradient 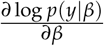 and the Hessian 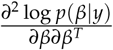 efficiently. Villani et al.(2012) show that this can be done very efficiently using the chain rule and compact matrix computations, and Appendix A gives the details for the NC-*χ* regression. In DTI, when the parameter space is restricted to the set of positive definite matrices, these expressions need to be extended, see Section 5.2.

In summary, our proposed algorithm consists of a two-block Metropolis-Hastings within Gibbs sampler, where each updating step updates a set of regression coefficients simultaneously with their binary variable selection indicators. The multivariate student-*t* proposal is tailored to the full conditional posterior at each step, using a Newton method to approximate the conditional posterior mode and curvature (Hessian). The computations are very fast since the gradient and the Hessian for the Newton steps can be computed very efficiently in compact matrix form, and only a very small number of Newton steps is needed, since each iteration starts at the previously accepted parameter draw which is typically an excellent initial value.

## 4. Activity localization in FMRI data

Comparisons of the proposed Rician model (Eq. 2.1) to a corresponding Gaussian model using several commonly used fMRI datasets showed no detectable differences between the two models since the SNRs were larger than three in all voxels; this is in line with the results in (Rowe and Logan, 2004). As discussed in the Introduction, however, there are situations when SNRs can be low in fMRI, in particular for high-resolution imaging. We therefore compare the two models using simulated fMRI data with Rician noise at the three different SNR levels (1, 2 and 3). The data are simulated from a model that mimic the results from a real fMRI experiment with a simple block paradigm. The real fMRI data had a spatial resolution of 1.6 x 1.6 x 1.8 mm^3^, and the noise variance and the variance for the activation parameter was manipulated in the simulated datasets to obtain a pre-specified level of activation and SNR. The noise variance experimentally controls the SNR levels, while the variance for the activation parameter is adjusted to accommodate one of the four chosen *t*-ratios (0, 3, 5 and 7) for each of the voxels on our selected slice of the brain. The prior distributions on the parameters in the Rician and Gaussian model are carefully chosen to carry the same information in both models. Specifically, we choose unit information priors (see Section 3.2), such that the priors only carry the information from a single observation in each of the models. We simulate 100 datasets for each SNR level (1, 2, 3). The first row with graphs in Figures 4.1 and 4.2 shows the four activated regions in the data generating process in the form of “*t*-ratios” (parameter value/standard deviation). The second row of graphs show that all four activation regions are correctly localized with our Rician model in a large majority of the simulated datasets, while the third row shows that the Gaussian model completely misses all of the activated regions when SNR=1, and has a high failure rate when SNR=2.

**FIGURE 4.1.**
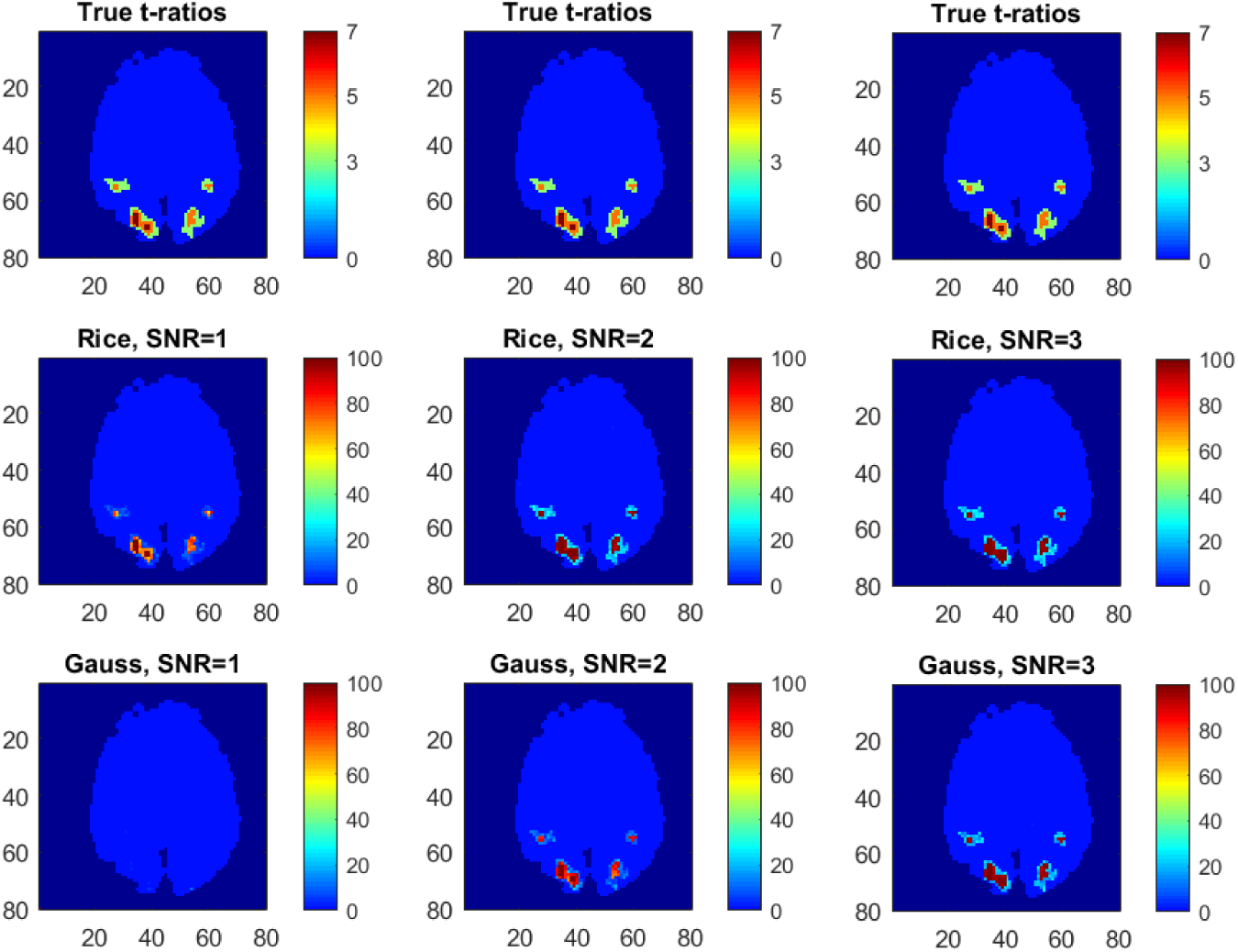
Comparison of activation inferences using the Rician and Gaussian models on simulated fMRI data. Top row: True activations in the Rician data generating model. Middle (Rice) and bottom (Gauss) row: Percentage of simulated datasets where the posterior probability of activation is larger than 99 %. Color should be used in print.

**FIGURE 4.2.**
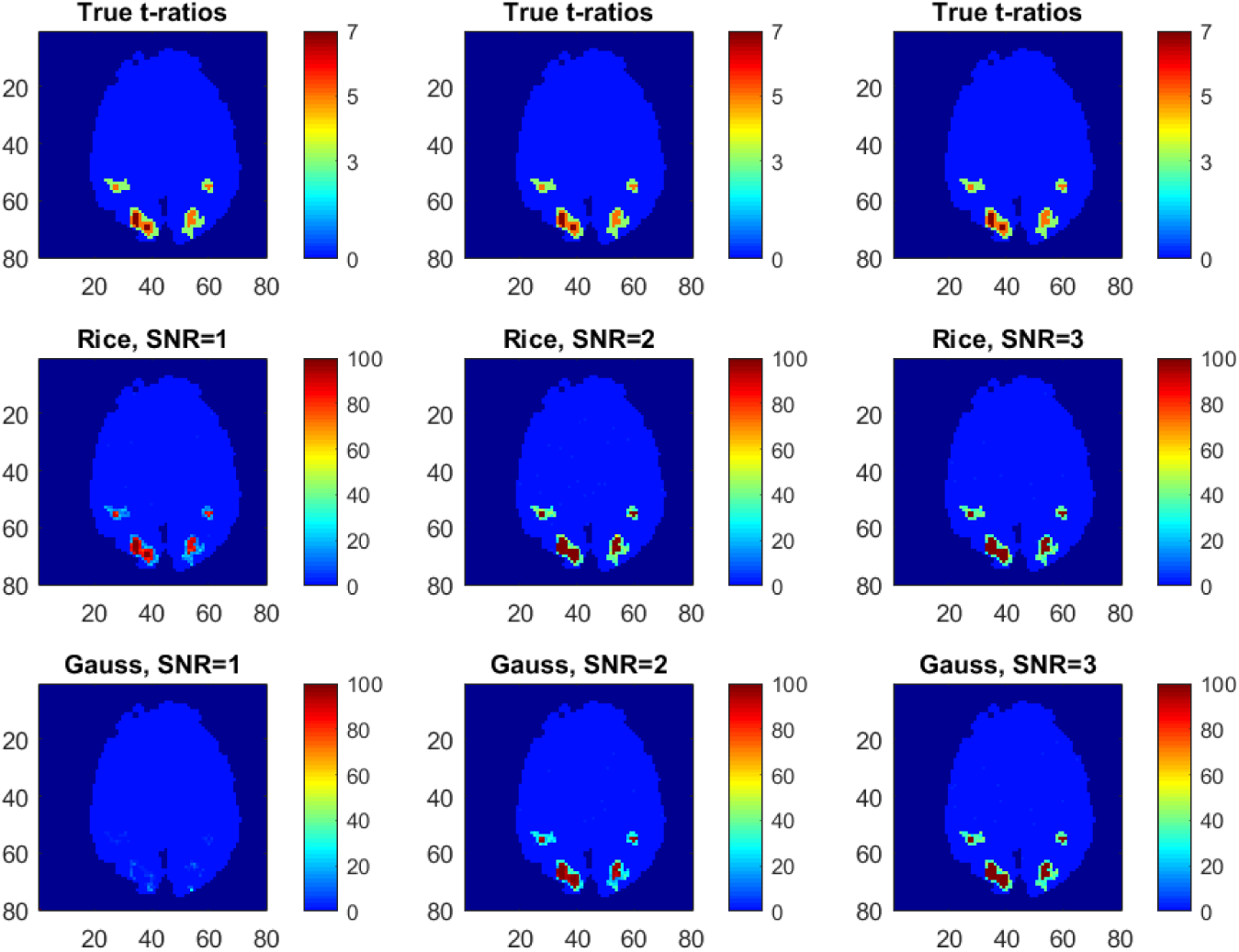
Comparison of activation inferences in the Rician and Gaussian models using simulated fMRI data. Top row: True activations in the Rician data generating model. Middle (Rice) and bottom (Gauss) row: Percentage of simulated datasets where the posterior probability of activation is larger than 95 %. Color should be used in print.

## 5. Estimating fractional anisotropy and mean diffusivity in DWI data

### 5.1. Diffusion weighted imaging

While fMRI data are mainly specified by the echo time and the repetition time of the pulse sequence, DWI data also require specification of the *b*-value (Le Bihan et al., 1986). The *b*-value in turn depends on two factors; the strength and the duration of the diffusion gradient. Using a larger *b*-value enables more advanced diffusion models, e.g. through HARDI (Tuch et al., 2002), which for example can be used to properly account for multiple fiber orientations in a single voxel. A significant drawback of a higher *b*-value is, however, a lower signal to noise ratio. The main reason for this is that the signal decays exponentially with time, and high b-values require longer diffusion gradients. As a consequence, Rician noise models are far more common for DWI than for fMRI, as the Rician distribution is only well approximated by a Gaussian for high SNRs.

### 5.2. The diffusion tensor model

The most common diffusion tensor model states that the signal *S_i_* for measurement *_i_* can be written as

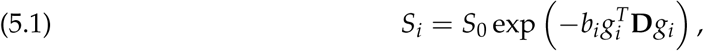

where *S*_0_ is the signal in absence of any diffusion gradient, *b_i_* is the *b*-value, g*_i_* = (g*_ix_*, *g_iy_*, g*_iz_*)*^T^* is the gradient vector and

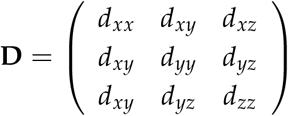

is the diffusion tensor. The single-diffusion tensor model in (5.1) can be written as a regression model of the form in Eq. (2.3) with (see e.g. Koay (2011))

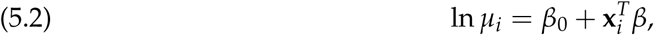

where *β*_0_ In *S*_0_, *β* (*d_xx_*, *d_yy_*, *d_zz_*, *d_xy_*, *d_yz_*, *d_xz_*) and

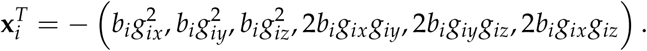

For single-coil imaging, the noise around *μ_i_* is Rician, and cannot be well approximated by a Gaussian model for high *b*-values where the signal-to-noise ratio is low. When data are collected by parallel techniques using *L* coils, the noise is either Rician distributed or NC-*χ* distributed with *L* degrees of freedom. If the composite signal is a complex weighted sum of the *L* signals, the magnitude of the composite signal is Rician distributed. If the simpler sum of squares approach is used for merging the L signals into a single image, the resulting signal is instead NC-*χ* distributed (Tristán-Vega et al., 2012; Aja-Fernandez and Vegas-Sanchez-Ferrero, 2016).

Note that since the tensor *D* is positive definite, the parameter space of *β* in (5.2) is restricted. One can impose the positive definitness restriction by assigning zero prior probability to all *β* that correspond to a negative definite *D*; all such proposals will then be rejected in the MCMC. This may, however, lead to excessive rejections, and a better solution is to impose the positive definiteness restriction explicitly. We here use the Log-Cholesky representation (Koay, 2011), where the diffusion tensor *D* is expressed as

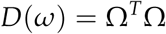

with

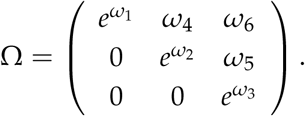

In this parametrization the tensor can be written as

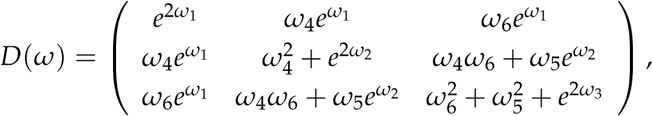

such that the vector of regression coefficients *β*(*ω*) is given by

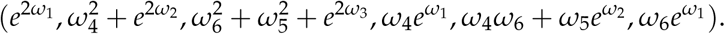

Most applications with the diffusion tensor model takes the logarithm of the measurements and estimates *β* with least squares (see Koay (2011) for an overview). This estimation method therefore does not respect the log link in the mean. One can also argue that it also implicitly assumes Gaussian noise in the sense that least squares equals the maximum likelihood estimate only when the noise is Gaussian. Moreover, it does not guarantee that the estimated tensor is positive definite. We refer to Koay (2011) for an overview of constrained non-linear least squares alternatives.

We will here take a Bayesian approach with Rician or NC-*χ* noise, using a proper log link and a parametrization that guarantees that the posterior mass is fully contained within the space of positive definite matrices. Existing Bayesian approaches to DTI assume Gaussian noise and use the random walk Metropolis (RWM) algorithm to simulate from the posterior distribution. RWM is easy to implement, but is well known to explore the posterior distribution very slowly (see Section 5.4). The Metropolis-within-Gibbs algorithm with tailored proposals and variable selection to reduce the dimensionality of the parameter space presented in Section 3.4 can explore the posterior distribution in a much more efficient manner (Villani et al., 2009, 2012). As a result of the non-linear mapping from *ω* to *β*, the gradient of the likelihood part of Equation A.2 is modified to

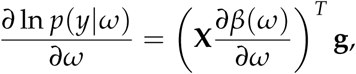

where

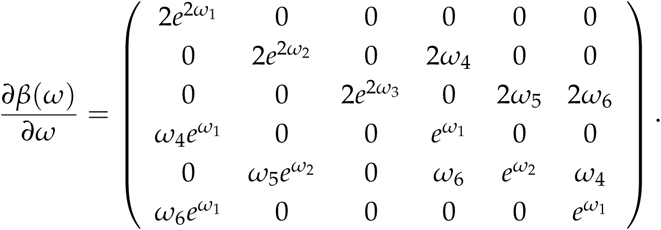

The Hessian in Equation A.3 can be modified accordingly.

The Fisher information based prior presented in Section 3.2 can in principle be used for DTI. We have found however that the numerical stability of our MCMC sampler improves if we use an alternative prior, which we now describe. We assume the priors for the intercepts *β*_0_ ~ *N*(*m_β_*,*d*) and *α*_0_ ~ *N*(*m_α_*,*d*), independently of the priors for the unrestricted tensor coefficients *ω* ~ *N*(0,*cI*) and the parameters variance function *α* ~ *N*(0,*cI*), where *c* = 100 to induce non-informative priors and *I* is the identity matrix. Note that the prior expected value of 0 for *α* implies that the variance of the underlying complex-valued signal *ϕ* is centered on the homoscedastic model a priori. To set the prior mean on the intercepts *β*_0_ and *α*_0_, note first that the models for *μ* and *σ^2^* in Eq. 2.1 become *β*_0_ = ln *μ_i_* and *α*_0_ = ln *σ_i_* when *b* = 0. It is therefore common in the DTI literature to separately pre-estimate the mean intercept *β*_0_ by the logarithm of the mean of measurements *y* when *b* = 0, and then subsequently remove these observations from the dataset. This procedure improves the numerical stability of the estimations. In a similar vein, we set the prior expected values, *m_β_* and *m_α_* by taking the logarithm of the mean and variance of *y* when *b* = 0, respectively; the observations with zero *b*-values are then removed from the dataset in the remaining estimation. We have found improved numerical stability in the MCMC algorithm if we allow for a positive, but small, prior variance of *d* = 0.1.

## 5.3. Data

We use the freely available MGH adult diffusion dataset from the Human Connectome Project (HCP) (Setsompop et al., 2013; Essen et al., 2013) ^1^. The dataset comprise DWI data collected with several different *b*-values, and the downloaded data have already been corrected for gradient nonlinearities, subject motion and eddy currents (Glasser et al., 2013; Andersson and Sotiropoulos, 2016). The DWI data were collected using a spin-echo EPI sequence and a 64-channel array coil (Setsompop et al., 2013), yielding volumes of 140 x 140 x 96 voxels with an isotropic voxel size of 1.5 mm. The data collection was divided into 5 runs, giving data with four different *b*-values: 1,000, 3,000, 5,000, and 10,000 s/mm^2^. The number of gradient directions was 64 for *b* = 1,000 s/mm^2^ and *b* = 3,000 s/mm^2^,128 for *b* = 5,000 s/mm^2^, and 256 for *b* = 10,000 s/mm^2^. Merging the measurements from the 64 channels into a single image was performed using a complex weighted combination (Setsompop et al., 2013), instead of the more simple sum of squares approach. This is an important fact, as the weighted approach for this data leads to noise with a Rician distribution, instead of the NC-*χ* distribution resulting from the sum of squares approach (Aja-Fernandez and Vegas-Sanchez-Ferrero, 2016). Prior to any statistical analysis, the function FAST (Zhang et al., 2001) in FSL was used to generate a mask of white brain matter, gray brain matter and cerebrospinal fluid (CSF), to avoid running the analysis on voxels in CSF.

Data used in the preparation of this work were obtained from the Human Connectome Project (HCP) database (https://ida.loni.usc.edu/login.jsp). The HCP project (Principal Investigators: Bruce Rosen, M.D., Ph.D., Martinos Center at Massachusetts General Hospital; Arthur W. Toga, Ph.D., University of Southern California, Van J. Weeden, MD, Martinos Center at Massachusetts General Hospital) is supported by the National Institute of Dental and Craniofacial Research (NIDCR), the National Institute of Mental Health (NIMH) and the National Institute of Neurological Disorders and Stroke (NINDS). HCP is the result of efforts of co-investigators from the University of Southern California, Martinos Center for Biomedical Imaging at Massachusetts General Hospital (MGH), Washington University, and the University of Minnesota.

## 5.4. Comparisons between the Rician and Gaussian DTI models

We compare the Rician and Gaussian DTI models for the voxels in slice 50 in the middle of the brain. We mainly compare the estimation results between the models using the whole dataset with all b-values up to *b* = 10,000 s/mm^2^, but also show some results for subsets of the whole dataset with b-values up to *b* = 3,000 s/mm^2^ and *b* = 5,000 s/mm^2^, respectively. The expected Hessian is used for the MH proposals of the parameters in the Gaussian case, but since the expected Hessian is not available for the Rician model, different combinations of the observed Hessian and the outer product of gradients for *μ* and *ϕ* are used in each voxel, to improve the numerical stability of the estimations. Our MCMC convergence is excellent for both the Rician and Gaussian DTI models, with high acceptance probabilities for *μ* and *ϕ* in almost all voxels for all estimated datasets. The mean MH acceptance probabilities for *μ* and *ϕ* are 74 % and 87 % for the Rician model, compared to 70 % and 90 % for the Gaussian model. The standard deviations of the acceptance probabilities across voxels are 7.5 % and 16.2 % for the Rician model, compared to 5.1 % and 5.2 % for the Gaussian model.

We compare the efficiency of our MCMC algorithm to commonly used Random Walk Metropolis (RWM) algorithms for MCMC in DTI (see e.g. the highly influential work by Behrens et al. (2003)). The RWM algorithms use a multivariate normal distribution centered on the current parameter value to propose a posterior draw of all parameters in *μ* and *σ* in a single block. The most common choice of proposal covariance matrix in DTI is a scaled identity matrix where the scale is chosen adaptively to achieve optimal performance. We also compare our MCMC algorithm to a refined version with covariance matrix —*cH*^−1^, where *H* is the Hessian at the posterior mode and *c* is a scalar which is again chosen adaptively for optimal performance. Using the posterior results from 100 randomly sampled white matter voxels, Figures 5.1 and 5.2 show histograms of the ratios of the number of independent draws per minute for our MCMC algorithm compared to each type of RWM algorithm. Results are presented for both the Rician and Gaussian models. The number of independent MCMC draws is defined as the number of total MCMC draws divided by the estimated inefficiency factor 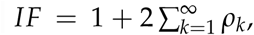, where *ρ_k_* is the autocorrelation function of lag *k* of the MCMC chain.

In general, our MCMC algorithm is much more efficient in almost all voxels than the RWM algorithm with covariance matrix *cI* for both the Rician and Gaussian models. In most voxels, our MCMC algorithm for the Gaussian model is also more efficient than the RWM algorithm with covariance matrix —*cH*^−1^. This is also true for our MCMC algorithm in a majority of the voxels for the Rician model, especially for parameters *β_μ_* compared to *β_σ_*. In a random sample of 100 gray matter voxels, we also find that our MCMC algorithm is much more efficient than the RWM algorithm with covariance matrix *cI*, but compared to the RWM algorithm with covariance matrix −*cH*^−1^ our algorithm is only slightly better in both models (not shown here).

**FIGURE 5.1.**
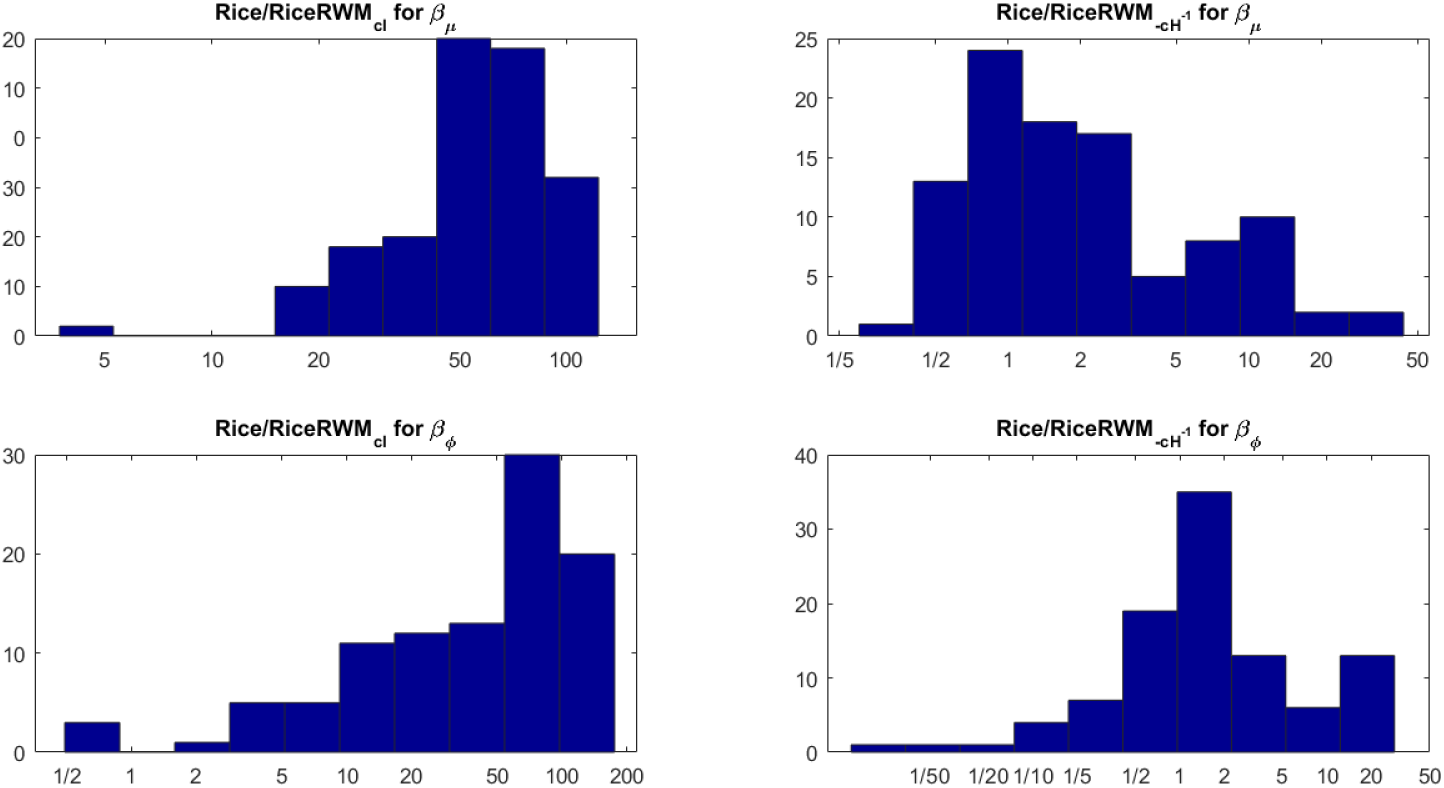
Histograms of the ratio of independent draws per minute for our MCMC algorithm compared to each type of RWM algorithm for 100 randomly sampled white matter voxels for the Rician model. The rows correspond to the parameters, the columns to the two covariance matrices in the RWM algorithm.

**FIGURE 5.2.**
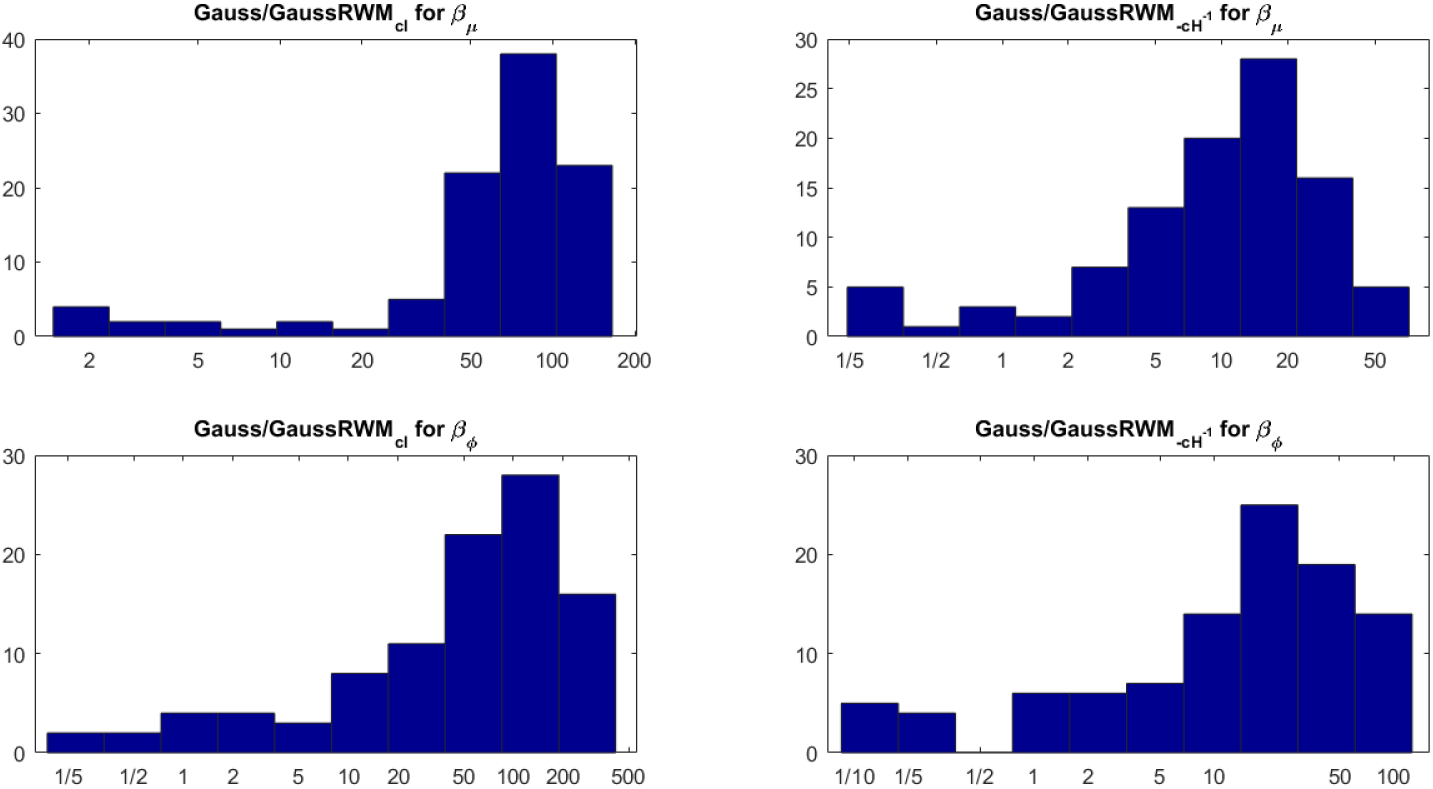
Histograms of the ratio of independent draws per minute for our MCMC algorithm compared to each type of RWM algorithm for 100 randomly sampled white matter voxels for the Gaussian model. The rows correspond to the parameters, the columns to the two covariance matrices in the RWM algorithm.

Figure 5.3 shows posterior inclusion probabilities for the covariates corresponding to (*d_xx_*, *d_yy_*, *d_zz_*) in **z** (the variance function) for both models. In a large number of voxels the inclusion probabilities for the Gaussian model are close or equal to 1, compared to far fewer voxels for the Rician model. In addition, there are substantially more voxels with this property in the mid-regions of the brain for the Gaussian model, and in the outer parts of the brain for the Rician model. The inclusion probabilities for the remaining covariates in z are in most cases very close to zero for both models, especially for the Gaussian model (not shown here). This clearly shows that diffusion covariates affect the noise variance in both models, and may imply that homoscedastic DTI models can give distorted results as documented in Wegmann et al. (2016) for the Gaussian DTI model. Using a part of the dataset with all b-values up to *b* = 5,000 s/mm^2^ (thereby excluding relatively uncommon measurements at a b-value of 10,000) implies far fewer voxels with inclusion probabilities close or equal to 1 for both models, but there are still substantially more voxels with this property for the Gaussian model compared to the Rician model (not shown here).

**Figure 5.3.**
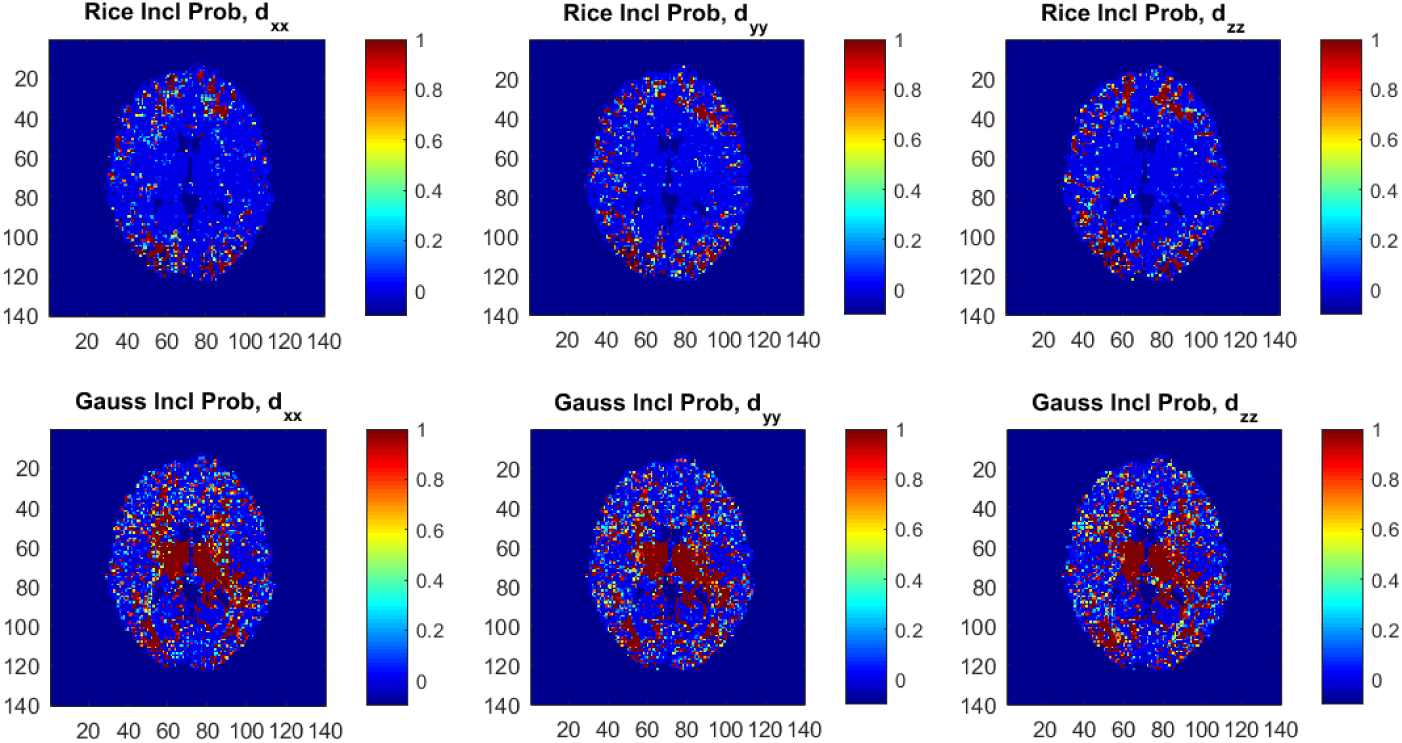
Posterior inclusion probabilities for the covariates corresponding to the diffusion directions (*d_xx_*, *d_yy_*, *d_zz_*) in the variance function *ϕ* for the Rician and Gaussian DTI models. Note that the inclusion probability is close to 1 for a large number of voxels, especially for the Gaussian model. Color should be used in print.

The estimated single-diffusion tensors are compared across voxels for the Rician DTI model in Eq. 2.1 to the Gaussian counterpart, with respect to the DTI scalar measures mean diffusivity (MD) and fractional anisotropy (FA). The DTI scalar measures are functions of the eigenvalues λ_1_ ≥ λ_2_ ≥ λ_3_ of the single-diffusion tensor, defined as

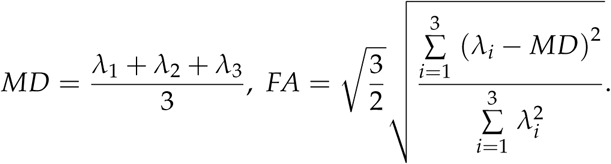

**FIGURE 5.4.**
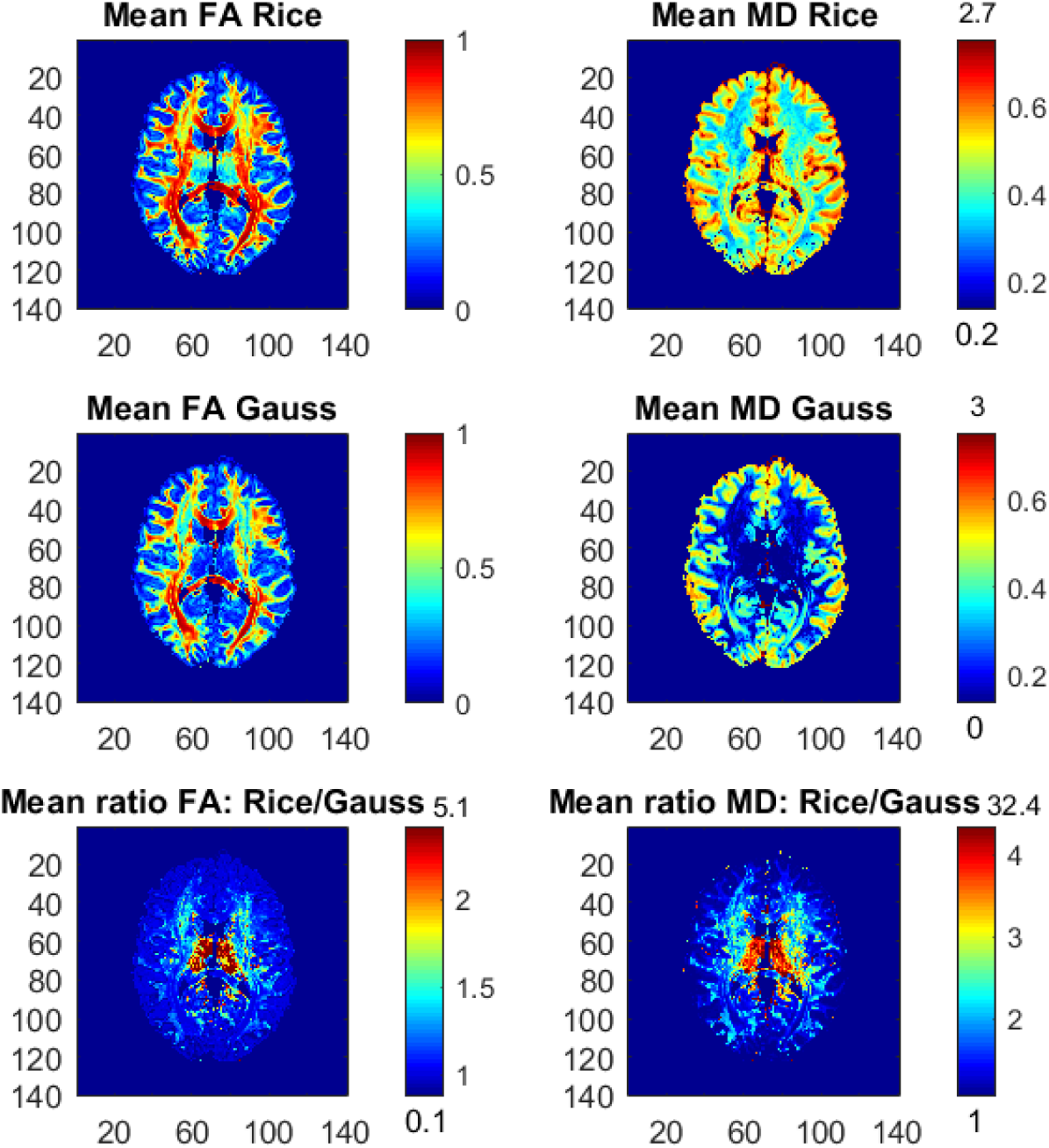
Posterior means and ratios of posterior means of FA and MD for the Rician and Gaussian DTI models, using the whole dataset with all b-values up to *b* = 10,000 s/mm^2^. The colorbars are shown for the mid 95 % values and the minimum and maximum values are marked out at the bottom and top of the colorbars, respectively. Color should be used in print.

Figure 5.4 shows the posterior means of FA and MD and the ratios of posterior means between the models, and Figure 5.5 shows the posterior standard deviations of FA and MD and the ratios of posterior standard deviations between the models.

In general, the Gaussian model substantially underestimates mean values of FA in many voxels, especially in mid-regions with low or mid-size values of FA, compared to the theoretically correct Rician model. In addition, the Gaussian model greatly underestimates MD across the whole slice of the brain compared to the Rician model. Hence, using the Gaussian model for DTI can therefore lead to severely misleading inferences. The standard deviations of FA and MD are small for both models. In white-matter regions with high FA values the Gaussian model estimates slightly larger standard deviations of FA compared to mid-regions with slightly larger standard deviations of FA for the Rician model. On the other hand, the standard deviations of MD are underestimated by the Gaussian model in all voxels.

Figure 5.6 shows the posterior means of FA and MD and the ratios of posterior means for the Rician models with covariates in the noise variance *ϕ* (heteroscedastic model) and without covariates in *ϕ* (homoscedastic model). The differences between the models are small, but in the outer parts of the brain the homoscedastic Rician model slightly overestimates the posterior means of FA in a large number of voxels. The posterior standard deviations of FA and MD for the Rician models are similar, but the homoscedastic Rician model slightly underestimates, in general, the standard deviation of FA in the outer parts of the brain (not shown here). The differences in FA between the Rician models agree with our previous findings that the diffusion covariates (directions) especially affect the noise variance for the Rician model in the outer parts of the brain, where directional DTI measures such as FA are affected. This is in contrast to the non-directional measure MD, for which the differences between the models are negligible. Hence, in voxels with heteroscedastic noise variance that depends on the diffusion directions the posterior means and standard deviations of FA are slightly different for the heteroscedastic and homoscedastic Rician models.

**Figure 5.5.**
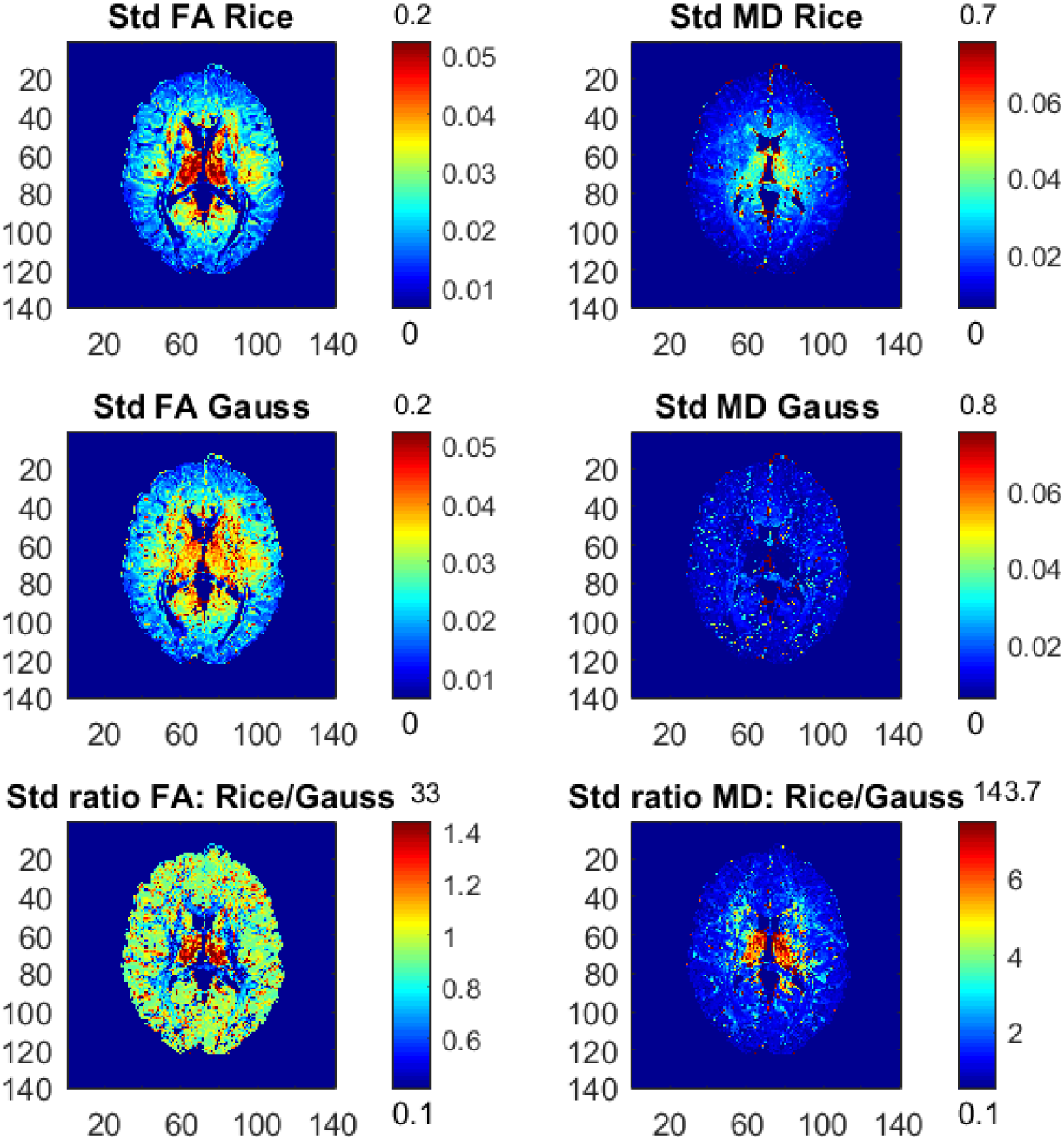
Posterior standard deviations and ratios of posterior standard deviations of FA and MD for the Rician and Gaussian DTI models, using the whole dataset with all b-values up to *b* = 10,000 s/mm^2^. The colorbars are shown for the mid 95 % values and the minimum and maximum values are marked out at the bottom and top of the colorbars, respectively. Color should be used in print.

It is relatively uncommon with measurements at a b-value of 10,000. Figure 5.7 therefore shows the posterior means and the ratios of posterior means of FA and MD between the models for the part of the whole dataset with all b-values up to *b* = 5,000 s/mm^2^, hence excluding the observations with the highest b-value. The differences in FA and MD are notably smaller compared to the results from the whole dataset, but the Gaussian model still underestimates the posterior mean values of FA and MD substantially in many voxels. Hence, using the Gaussian model can also lead to misleading inferences for this smaller subset of the data. Taking an even smaller data subset with all b-values up to *b* = 3,000 s/mm^2^, the differences in FA and MD between the models become negligible, where the Gaussian model only slightly underestimates FA and MD in some voxels (not shown here).

**FIGURE 5.6.**
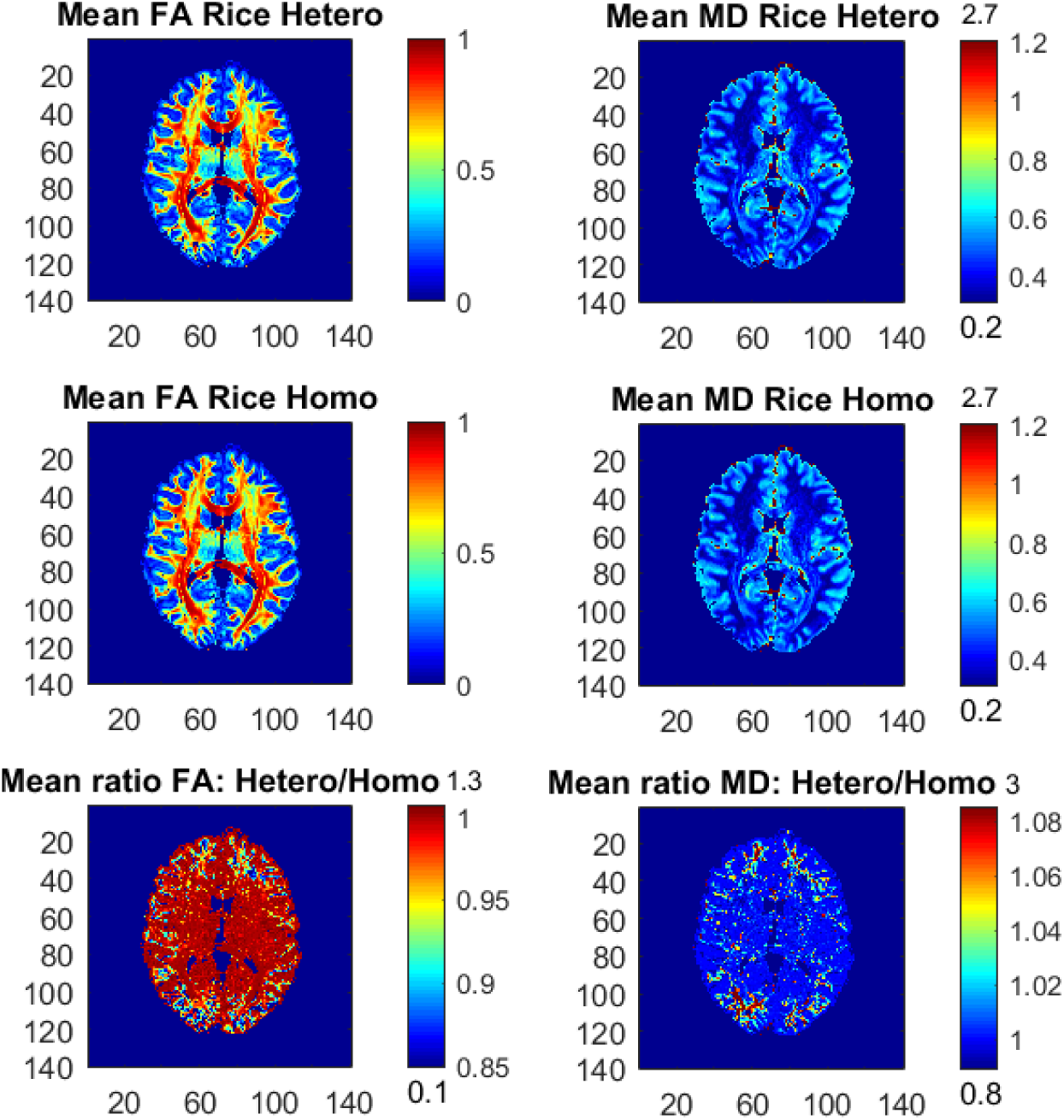
Posterior means and ratios of posterior means of FA and MD for the heteroscedastic (Hetero) and homoscedastic (Homo) Rician DTI models, using the whole dataset with all b-values up to *b* = 10,000 s/mm^2^. The colorbars are shown for the mid 95 % values and the minimum and maximum values are marked out at the bottom and top of the colorbars, respectively. Color should be used in print.

To investigate the differences between the Rician and Gaussian models in white and gray matter, we use the function FAST in FSL to compute the probabilities for white matter, gray matter and CSF in each voxel of the brain. Let a white-matter (gray-matter) voxel be defined as a voxel where the probability is 1 for white matter (gray matter). It is generally expected that white-matter voxels have higher FA, compared to gray-matter voxels. Figure 5.8 shows that this is true for the Rician model as the distribution of the posterior means of FA is more skewed to larger values, compared to more uniformly distributed posterior means of FA for the Gaussian model. Hence, the Gaussian model underestimates, on average, FA in white-matter voxels. In addition, Figure 5.8 shows that the uncertainty of FA for white-matter voxels is somewhat lower for the Rician model compared to the Gaussian model, with the distribution of the standard deviations of FA being more skewed to the left for the Rician model. This is in contrast to the gray-matter voxels, where the distributions of both the posterior means and standard deviations of FA are very similar and concentrated at low values for both models.

**FIGURE 5.7.**
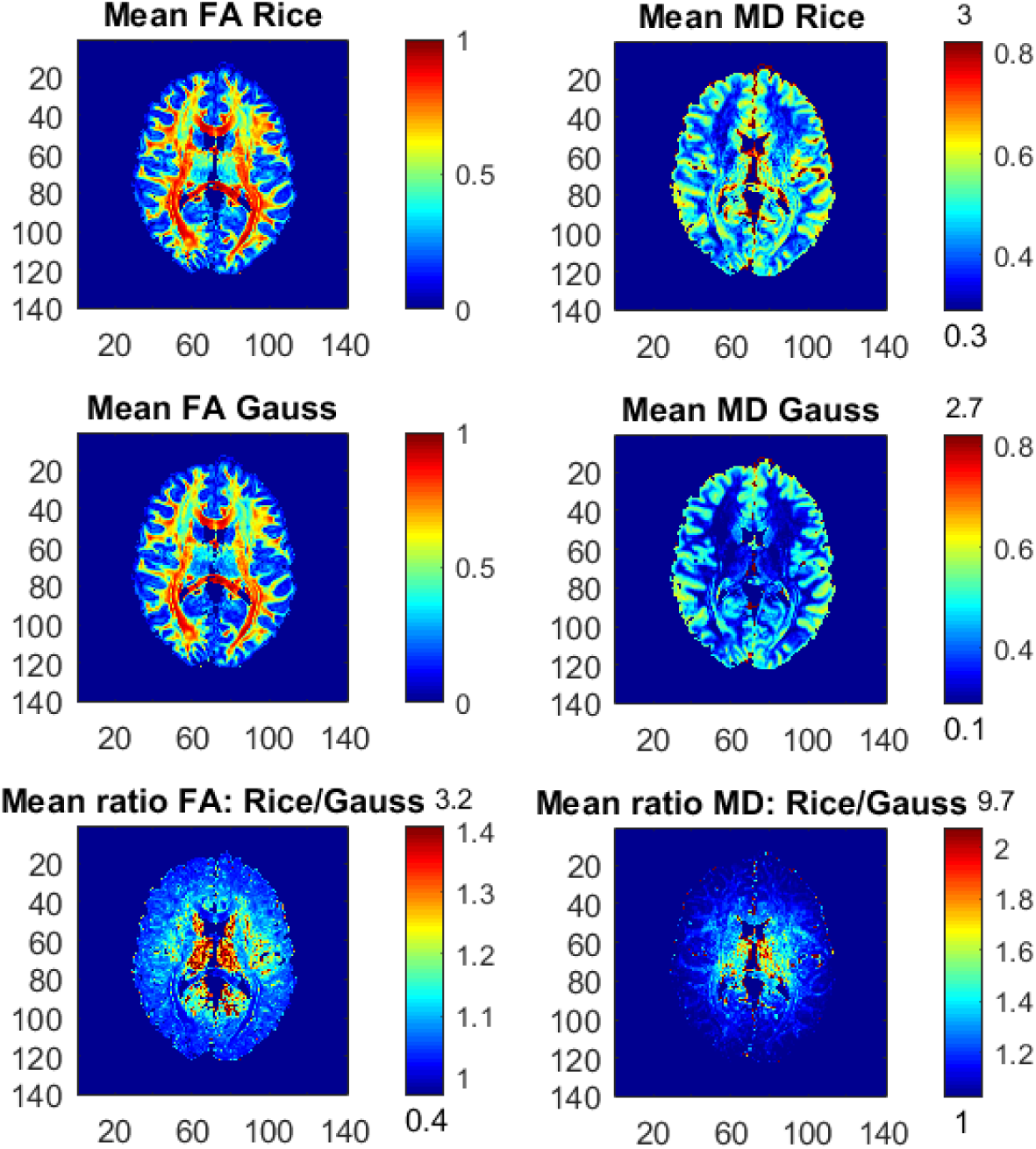
Posterior means and ratios of posterior means of FA and MD for the Rician and Gaussian DTI models, using the part of the dataset with all *b*-values up to *b* = 5,000 s/mm^2^. The colorbars are shown for the mid 95 % values and the minimum and maximum values are marked out at the bottom and top of the colorbars, respectively. Color should be used in print.

**FIGURE 5.8.**
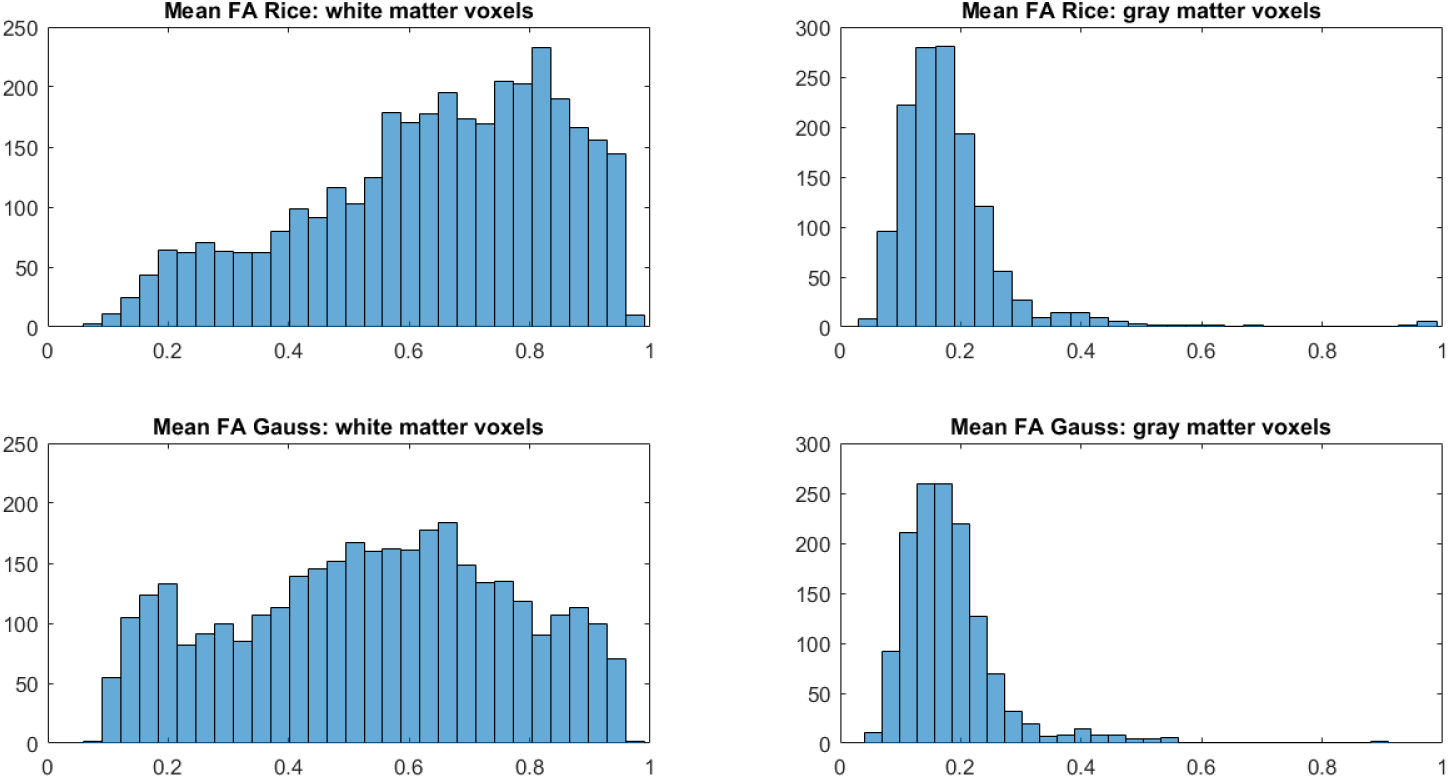
Histograms of posterior means of FA for white-matter (left) and gray-matter (right) voxels for the Rician and Gaussian DTI models, using the whole dataset. A white-matter (gray-matter) voxel is defined as a voxel where the probability is 1 for white matter (gray matter) from the function FAST in FSL.

**FIGURE 5.9.**
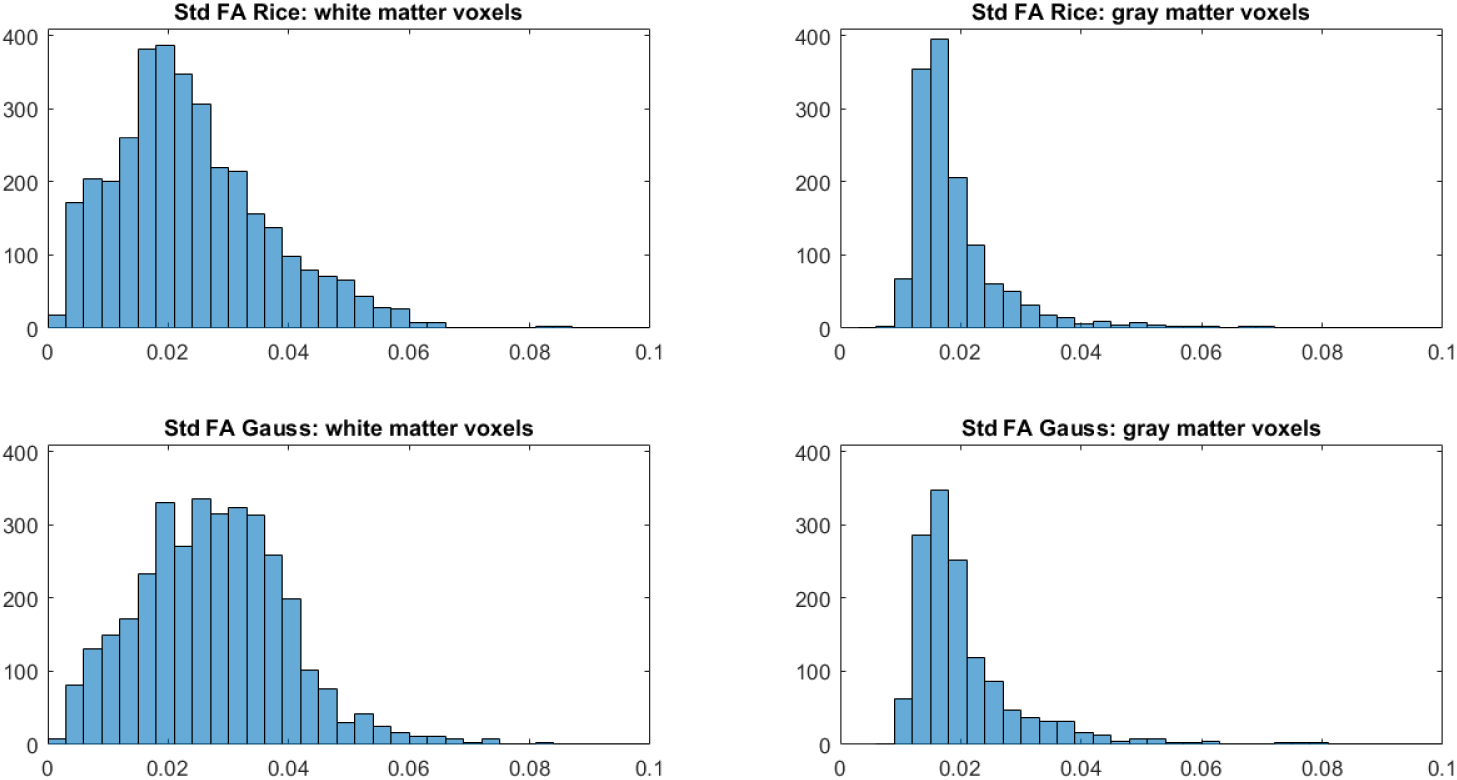
Histograms of posterior standard deviations of FA for white-matter (left) and gray-matter (right) voxels for the Rician and Gaussian DTI models, using the whole dataset. A white-matter (gray-matter) voxel is defined as a voxel where the probability is 1 for white matter (gray matter) from the function FAST in FSL.

## 6. Discussion

We propose a Bayesian non-central *χ* regression model for neuroimaging with the Rician model as a prominent special case. The model is applied to real diffusion data from the Human Connectome Project (Essen et al., 2013) and to simulated fMRI data with different SNRs. We show that the results from the theoretically correct Rician DTI model can differ substantially from the approximate Gaussian model typically used for diffusion tensor estimation. The Gaussian model greatly underestimates the mean diffusivity (MD) and substantially underestimates the FA of the single-diffusion tensors, which is consistent with previous results (Andersson, 2008). We also show that the differences between the Rician and Gaussian models increase with the b-value, which is natural since the SNR decreases with a higher b-value. Our results for real fMRI datasets are consistent with previous work (Solo and Noh, 2007; Adrian et al., 2013), which also come to the conclusion that there are negligible differences between the Rician and Gaussian noise models. We demonstrate, however, that the Rician model is remarkably adept at recovering the activations for simulated fMRI datasets at very low SNRs, which are more common in high-resolution images; we also show that the Gaussian model fails to detect activity for low SNRs.

Our framework is more general compared to the work by Andersson (2008) and other frameworks, as it is possible to include covariates for both the mean and the variance of the noise, and not only covariates for the mean. We show that DTI noise of the underlying complex-valued signal is heteroscedastic, especially for the Gaussian model. This is consistent to our recent work (Wegmann et al., 2016), where we showed that using diffusion covariates for the noise variance gives rather different results for DTI. It is also possible to include head motion parameters, and their temporal derivatives, as covariates for the noise variance for both fMRI and DTI. This can for example be used to down-weight measurements close to motion spikes (Power et al., 2014; Elhabian et al., 2014; Siegel et al., 2014) (as any measurement with a high variance is automatically down-weighted in our framework). For models with a large number of covariates, our variable selection can automatically discard covariates of no interest.

A potential drawback of our approach is the computational complexity. It takes 5.6 seconds to run 1,000 MCMC iterations for the Gaussian model in a representative voxel for the DTI data, and 11.2 seconds for the Rician model. For a typical DTI dataset with 20,000 brain voxels, this gives a total processing time of 31.1 hours for the Gaussian model and 62.2 hours for the Rician model. For this reason, we have only analyzed a single subject, as a group analysis with 20 subjects would be rather time consuming. As each voxel is analyzed independently, it is in theory straightforward to run MCMC on the voxels in parallel, using a CPU or a GPU (Eklund et al., 2013; Guo, 2012).

We have focused on the rather simple single-diffusion tensor, while more recent work focus on extending the diffusion tensor to higher orders. In the work by Westin et al. (2016), a regression approach is used to estimate the diffusion tensor and a fourth order covariance matrix in every voxel. Our regression framework can therefore easily be applied to QTI (q-space trajectory imaging) data (Westin et al., 2016) as well, and more generally for any diffusion model that can be estimated using regression. As a fourth order covariance matrix contains 21 independent variables, the possibility to perform variable selection becomes even more important. Furthermore, DTI is still the most common choice for studies investigating FA differences between healthy controls and subjects with some disease (Shenton et al., 2012; Eierud et al., 2014). Another indicator of the importance of FA is that the TBSS approach (Smith et al., 2006) has received more than 2,800 citations (with about 500 citations in 2015). Our approach gives the full posterior distribution of the FA, and any other function of the diffusion tensor, which can be used for tractography and to down-weight subjects with a higher uncertainty in a group analysis. This is in contrast to TBSS and the work by Andersson (2008), which ignore the uncertainty of the FA. Andersson (2008) develops a sophisticated maximum a posteriori (MAP) estimation for the DTI model, but does not deal with posterior uncertainty, in contrast to our full MCMC sampling from the posterior distribution.

## Acknowledgement

Anders Eklund was supported by the Information Technology for European Advancement (ITEA) 3 Project BENEFIT (better effectiveness and efficiency by measuring and modelling of interventional therapy) and by the Swedish research council (grant 2015-05356, “Learning of sets of diffusion MRI sequences for optimal imaging of micro structures“). Anders Eklund and Bertil Wegmann were supported by the Swedish research council (grant 2013-5229, “Statistical analysis of fMRI data”).

Data collection and sharing for this project was provided by the Human Connectome Project (HCP; Principal Investigators: Bruce Rosen, M.D., Ph.D., Arthur W. Toga, Ph.D., Van J. Weeden, MD). HCP funding was provided by the National Institute of Dental and Craniofacial Research (NIDCR), the National Institute of Mental Health (NIMH), and the National Institute of Neurological Disorders and Stroke (NINDS). HCP data are disseminated by the Laboratory of Neuro Imaging at the University of Southern California.

## Appendix A. Gradients and hessians

Villani et al. (2012) derive the gradient and the Hessian for a general posterior of the form

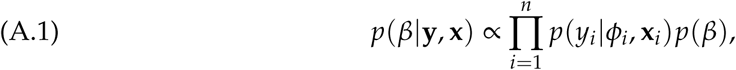

where 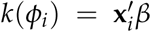 is a smooth link function, and **x***_i_* is a covariate vector for the *i*th observation; the full conditional posteriors for *β* and *α* in the Rician and the NC-*χ* case with logarithmic links are clearly of this form. The gradient of the likelihood in Eq. (A.1) can be expressed as

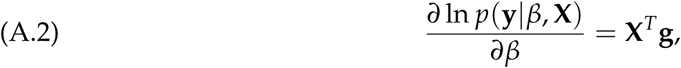

where **X** = (**x**_1_,…, **x***_n_*)*^T^*, **g** = (g_1_,…,gn)′,and

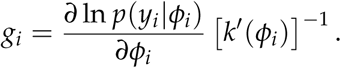

The Hessian of the likelihood is

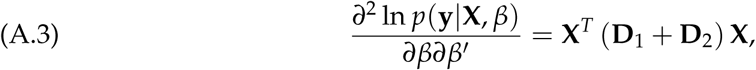

where **D**1 = Diag(*d*_1_*i*), **D**2 = Diag(*d*_2_*i*),

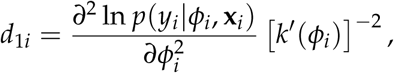

and

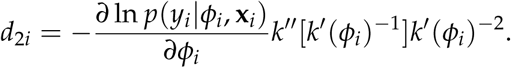

The outer-product approximation of the Hessian is given by

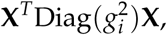

which is faster to compute and often numerically more stable than the Hessian itself. Finally, the Fisher information is given by

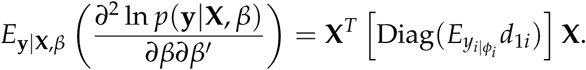

From (A.2) and (A.3) it is sufficient for our MCMC algorithm to compute the first and second derivatives with respect to (w.r.t.) *μ* and *ϕ*, respectively, for each of the individual observation. The log-likelihood for one observation *y* of a non-central *χ* variable can be written as (Tristán-Vega et al., 2012)

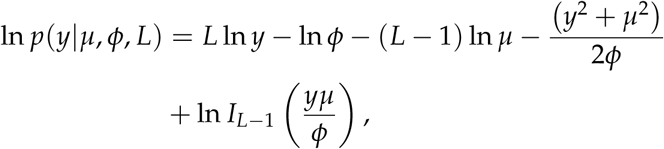
where *I_L_*—1(·) is the modified Bessel function of the first kind with order *L* — 1. The following derivatives hold for *I_L_*(*z*)

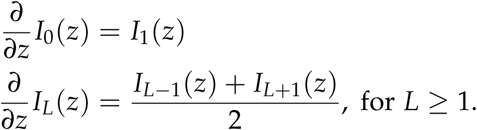

Let 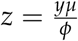 and

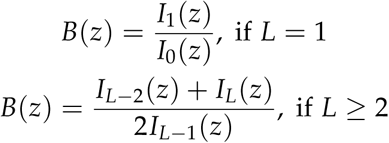

Then,

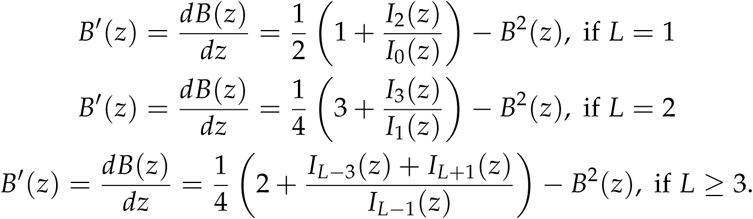

A.1. **Derivatives w.r.t. to** *μ*:

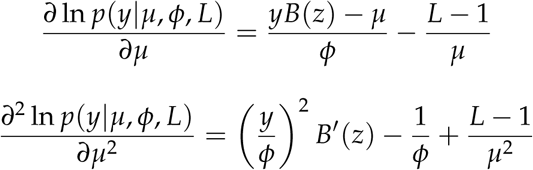

A.2. **Derivatives w.r.t. to** *ϕ*:

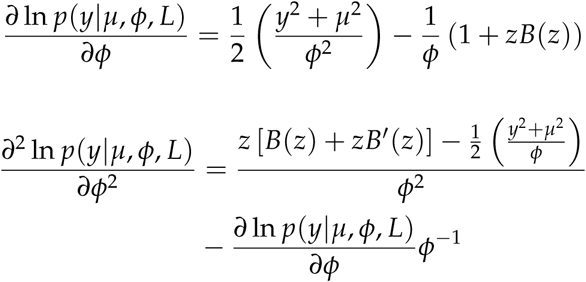

